# edgeR v4: powerful differential analysis of sequencing data with expanded functionality and improved support for small counts and larger datasets

**DOI:** 10.1101/2024.01.21.576131

**Authors:** Yunshun Chen, Lizhong Chen, Aaron T. L. Lun, Pedro L. Baldoni, Gordon K. Smyth

**Affiliations:** Bioinformatics Division, WEHI, Parkville, VIC 3052, Australia; ACRF Cancer Biology and Stem Cells Division, WEHI, Parkville, VIC 3052, Australia; Department of Medical Biology, The University of Melbourne, Parkville, VIC 3010, Australia; Computational Sciences, Genentech Inc, 1 DNA Way, South San Francisco, CA 94080, USA; School of Mathematics and Statistics, The University of Melbourne, Parkville, VIC 3010, Australia

## Abstract

edgeR is an R/Bioconductor software package for differential analyses of sequencing data in the form of read counts for genes or genomic features. Over the past 15 years, edgeR has been a popular choice for statistical analysis of data from sequencing technologies such as RNA-seq or ChIP-seq. edgeR pioneered the use of the negative binomial distribution to model read count data with replicates and the use of generalized linear models to analyse complex experimental designs. edgeR implements empirical Bayes moderation methods to allow reliable inference when the number of replicates is small. This article announces edgeR version 4, which includes new developments across a range of application areas. Infrastructure improvements include support for fractional counts, implementation of model fitting in C, and a new statistical treatment of the quasi-likelihood pipeline that improves accuracy for small counts. The revised package has new functionality for differential methylation analysis, differential transcript expression, differential transcript and exon usage, testing relative to a fold-change threshold and pathway analysis. This article reviews the statistical framework and computational implementation of edgeR, briefly summarizing all the existing features and functionalities but with special attention to new features and those that have not been described previously.

## Introduction

Next Generation Sequencing (NGS) has revolutionized biomedical research over the past 15–20 years. RNA-seq has become the standard technology for profiling gene and transcript expression [1, 2] while other technologies such as ChIP-seq, ATAC-seq, CUT&Tag, BS-seq and Hi-C allow high-resolution exploration of the molecular mechanisms by which expression is regulated [3].

edgeR is an R software package for differential analyses of data arising from NGS or similar technologies in the form of sequence read counts for genes or genomic features [4, 5, 6]. It is particularly designed to detect genes or features that have changed abundance levels between experimental conditions or cell types. edgeR pioneered the use of negative binomial (NB) generalized linear models (GLMs) to model read counts in genomic research [7, 8]. edgeR implements a range of novel statistical methods, including methods for borrowing information between genes, a strategy that is essential for genomic experiments with small sample sizes [4, 7, 9, 10]. It has become the underlying analysis engine for a wide range of sequencing technologies including ChIP-seq [11], Hi-C [12], bisulfite sequencing [13] and even proteomics [14, 15, 16]. The use of an explicit probabilistic count distribution allows edgeR to make meaningful inferences even for very low counts and provides a distinction with normal-based methods such as limma [17].

The edgeR package has undergone a number of major revisions since it was first released as part of the Bioconductor project in 2008 [18]. The original edgeR v1 pipeline (now called the “classic” pipeline) used exact conditional likelihood to achieve unbiased estimation of the NB-dispersion, exact NB tests to make pairwise comparisons between groups, and weighted likelihood empirical Bayes to borrow strength between genes [4, 5, 6]. These innovative statistical approaches allowed edgeR to achieve stable and reliable results even for experiments with very small numbers of biological replicates.

Full GLM functionality was introduced to edgeR in September 2010, allowing edgeR to model arbitrarily complex experiments including multiple treatment factors, batch effects and continuous covariates. All the original functionality was transferred to the GLM context, with Cox-Reid approximate conditional inference replacing the exact conditional likelihoods and likelihood ratio tests replacing the exact NB tests [7]. The edgeR GLM pipeline was released as edgeR v2 in 2011.

The second major revision was the introduction of quasi-likelihood (QL) methods in January 2012 [19, 20, 10, 21]. The QL model added a second dispersion parameter, the quasi-dispersion, which increased edgeR’s ability to model technical as well as biological sources of variability. Another key advantage was that the quasi-dispersions could be estimated by applying limma’s parametric empirical Bayes (EB) procedures to the genewise GLM deviances, which in turn enabled edgeR to leverage some of limma’s exact small-sample theory [22, 23]. The amount of EB moderation applied to the genewise dispersions could be optimized for the specific data at hand [9]. The GLM likelihood ratio tests could be replaced by quasi-F-tests, which allow for the uncertainty with which the genewise dispersions are estimated and thereby provide rigorous control of the false discovery rate (FDR) even for small sample sizes [19]. The edgeR QL pipeline was released as part of edgeR v3 in 2012.

This article announces edgeR version 4, which was released in October 2023 and includes new developments across a range of application areas. The revised package implements two fundamental changes that improve edgeR’s treatment of small counts and affect most analyses going forward. The first is a continuous generalization of the NB distribution that allows edgeR to accept fractional counts without rounding [24]. The second is a major revision of the QL pipeline based on improvements to the classical statistical theory underlying GLMs. The revision ensures unbiased quasi-dispersion estimates even when the read counts are very small and reduces computation time for large datasets with many samples [25]. Meanwhile, substantial parts of the edgeR package have been rewritten in C to increase speed and reduce memory usage.

The package also has new functionality for differential methylation analysis [13], differential transcript expression [24, 25], differential exon usage, differential transcript usage, testing relative to a foldchange threshold and pathway analysis.

This article reviews the statistical framework and computational implementation of edgeR, briefly summarizing all the existing features and functionalities but with special attention to the new features and those that have not been described previously.

## Materials & Methods

### Summary of the edgeR distributional model

#### Log-linear models

The input to edgeR is a matrix of sequence read counts with rows corresponding to genomic features and columns to biological samples. The rows often represent genes, but can correspond to almost any meaningful genomic feature including transcripts, exons, exon-exon junctions, genomic regions, methylation sites or DNA-DNA interactions.

Write *µ_gi_* for the expected number of reads assigned to feature *g* in sample *i*. edgeR assumes that *µ_gi_* can be modelled as a log-linear model

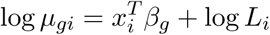

where *x_i_* is a covariate vector specifying the experimental conditions applied to sample *i*, *β_g_* is a coefficient vector that captures the experimental effects and log-fold-changes, and *L_i_* is the effective library size (sequencing depth) for sample *i*. From a mathematical point of view, the purpose of edgeR is to test hypotheses about *β_g_*.

#### Technical and biological variation produce a quadratic mean-variance relationship

Write *y_gi_* for the actual number of reads assigned to feature *g* in sample *i*. edgeR assumes that technical replicates produced from the same RNA or DNA sample (and with the same library size) would result in quasi-Poisson repeatability with variance 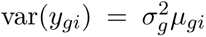. This represents the measurement error associated with sequencing and read alignment of a single sample. In ideal circumstances, the Poisson dispersion 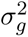 is close to 1 [26], but can be much greater than 1 in the presence of PCR duplication or if reads are probabilistically assigned to features [24, 25]. Singlecell data and transcript-level quantification are two application areas for which Poisson dispersions larger than 1 are common.

edgeR also assumes that the true abundance of feature *g* varies between biological replicates according to a gamma distribution with squared coefficient of variation equal to *ψ_g_*. The read count distribution therefore follows a mixture distribution across biological replicates with a quadratic meanvariance relationship of the form

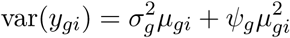

where 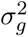 and *ψ_g_* represent technical and biological variation respectively [7, 24]. In edgeR terminology, 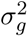 the quasi-dispersion and is the squared biological coefficient of variation (SBCV). The mean-variance relationship is usually rewritten as the variance function for a quasi-NB GLM,

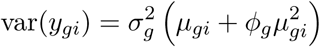

Where 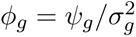 is the NB-dispersion parameter. If 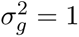, then the counts *y_gi_* can be considered to follow a NB distribution. If 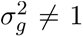, then *y_gi_* is distributed as a gamma mixture of quasi-Poisson distributions and quasi-NB methods are used.

#### Empirical Bayes

edgeR implements two empirical Bayes estimation strategies, a weighted likelihood empirical Bayes strategy for estimating the NB-dispersions, and a limma-style parametric empirical Bayes strategy for estimating the quasi-dispersions. The edgeR’s classic and GLM pipelines assume the quasi-dispersions are equal to 1 and use weighted likelihood empirical Bayes to estimate feature-specific NB-dispersions. The more recent QL edgeR pipelines set the NB-dispersions to global (trended or constant) values and then apply limma-style empirical Bayes to estimate feature-specific quasidispersions.

#### Divided counts

Baldoni *et al.* [24, 25] introduced the new concept of divided counts, which is relevant when the sequence reads have been probabilistically assigned to genes or transcripts, and technical resamples of the reads are available for each sample. In that context, the technical resamples can be used to estimate the part of the quasi-dispersion that arises from ambiguous assignment of reads to the genes or transcripts. The ambiguity dispersions can then be, literally, divided out of the read counts for each gene or transcript, leaving fractional counts that follow the same NB variance function but with reduced quasi-dispersion [24, 25]. The divided count strategy has the added advantage of strengthening the empirical Bayes moderation of the quasi-dispersions, because the ambiguity dispersions can vary by orders of magnitude and are a function of gene annotation topology, whereas the remaining quasi-dispersions are less heterogeneous and tend to be intensity-dependent. The divided counts have reduced count sizes compared to the original counts, reflecting the reduced information content that ambiguous read assignment represents.

#### Observation-level weights and library sizes

edgeR allows users to specify library sizes for individual observations if desired, which generalizes the log-linear model to

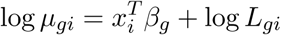

where *L_gi_* is the observation-specific effective library size. In edgeR terminology, the matrix of log *L_gi_* values is the offset matrix. Observation-specific library sizes can be used to implement non-linear normalization in edgeR [27, 28, 29, 12, 11] or to allow for transcript length effects [30]. edgeR v4 also allows observation-specific dispersions by

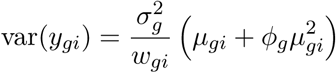

where the *w_gi_* are known GLM weights. The weights can be used to downweight outlier samples or observations [31].

### Example datasets

Example edgeR diagnostic plots are shown in Figure 3. The MDS plot in panel (a) is generated from the RNA-seq profiles of 882 human breast tumour samples from The Cancer Genome Atlas (TCGA) [32]. Each sample is classified as one of the five breast cancer intrinsic subtypes based on PAM50 gene signatures [33]. The data was downloaded from the TCGA Data Portal (https://tcga-data.nci.nih.gov) in the form of genewise read counts and processed as previously described [34]. Panels (b), (c) and (d) display a BCV plot, a quasi-dispersion plot, and a mean-difference (MD) plot respectively from the RNA-seq differential expression analysis described by Chen *et al.* [10]. The RNA-seq data for this analysis is from Fu *et al.* [35]. Panel (e) shows a differential splicing plot for the *Foxp1* gene. The *Foxp1* exons that are differentially used under the comparison are highlighted in the plot. The RNA-seq data for the differential exon usage analysis is from Fu *et al.* [36].

## Results

### Statistical principles

#### Differential analyses

edgeR is designed to conduct differential analyses of sequence read counts obtained from next generation sequencing (NGS) or similar technologies [6]. A dataset consists of a matrix of read counts where rows represent genomic features such as genes, exons, transcripts, genomic intervals, methylation sites or DNA-DNA interactions, and columns are samples associated with different biological groups or treatment conditions. In the following, we will refer to the genomic features as “genes”, with the understanding that the methodology is applicable to any genomic feature for which read counts can be obtained. The fact that edgeR focuses on differential results rather than on quantification of abundance means that edgeR can focus directly on statistical analysis of the read counts.

#### The negative binomial distribution captures biological variations

edgeR v1 and v2 assumed that the read counts follow NB distributions, an assumption that can be justified by a mixture model in which the true expression of each gene varies between biological replicates with constant coefficient of variation and the measurement error for individual samples follows a Poisson law [7, 24]. The use of an explicit count distribution allows meaningful probability calculations even for very small counts. The NB distribution implies a quadratic mean-variance relationship in which technical variation dominates for small counts and biological variation dominates for large counts. This assumption has been validated by extensive data analyses showing that NGS datasets do show the sort of quadratic mean-variance relationship that the NB distribution implies. The NB-dispersion parameter estimates the coefficient of variation of the true expression levels. The square-root of the NB-dispersion is called the biological coefficient of variation (BCV) in edgeR [7]. edgeR v3 generalized the NB assumption to allow quasi-dispersions and observation-specific weights, as described in Methods. If probability calculations are required and quasi-dispersions or weights are present, then the counts are interpreted as being averages of independent technical replicates. Using the same notation as in Methods, the conceptual number of technical replicates leading to count *y_gi_* is given by 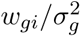.

#### Generalized linear models handle complex experiments

Another advantage of the NB distribution is that it belongs to the family of distributions for which generalized linear modelling is possible [8]. The NB GLM framework adopted in edgeR provides great flexibility for analysing complex multifactor experiments. Different experimental conditions and batch effects can be easily handled by the design matrix. It enables time-course analyses by incorporating a spline curve across different time points into the design [37]. Very general analyses are possible, for example gene-gene correlations can be detected by adding log-expression values for a target gene as a covariate column in the design matrix.

#### Unbiased estimation of the NB-dispersion

Obtaining an unbiased estimator of the NB-dispersion is not straightforward because regular maximum likelihood estimation is biased leading to under-estimated dispersions [5]. Unbiased estimation requires the likelihood be adjusted for estimation of the linear model parameters by conditioning on the linear model estimators, a process analogous to REML for normal-based models [38]. edgeR originally implemented an exact conditional likelihood approach, which was very effective but was applicable only to relatively simple oneway experiments with multiple groups but no blocks or covariates [5]. In edgeR v2, Cox-Reid adjusted profile likelihood (APL) was adopted under the more flexible GLM framework [39, 7]. This allows unbiased estimation of the NB-dispersions even for analyses with complex experimental designs [7].

#### Weighted likelihood empirical Bayes for the NB-dispersion

Simple genewise estimation of the NB-dispersion is not reliable for genomic datasets, which typically have many genes or genomic features but relatively few biological replicates. Given the parallel nature of genomic data, whereby the same log-linear model is fitted in parallel to a large number of genes, the accuracy of dispersion estimate for each individual gene can be greatly improved by adopting an empirical Bayes strategy to borrow strength between genes. A straightforward parametric empirical Bayes approach is not available for the NB-dispersion because the form of the probability distribution doesn’t make a conjugate prior distribution model possible. Instead, a weighted likelihood approach was implemented in edgeR to produce an approximate empirical Bayes strategy [4, 40]. The weighted likelihood approach has the advantage that it makes no assumptions about the shape of the prior distribution but instead adapts to the data at hand. The extension of the weighted likelihood approach to the NB GLM framework was described by McCarthy et al [7]. Under the GLM framework, a common NB-dispersion for all the genes can be estimated by maximizing the total APLs of all the genes with equal weights. To account for the fact that genes with lower expression level tend to have larger dispersions, trended NB-dispersions are introduced and estimated by maximizing the locally shared APL formed by genes with similar expression levels. Finally, a gene-specific NB-dispersion can be obtained by maximizing the weighted APLs formed by the individual APL of that gene and its locally shared APL with a proper weight. The weight represents the amount of prior information borrowed from the neighbouring genes. The final gene-specific NB-dispersion estimate can be considered as a compromise between the estimates obtained from the data for that gene alone and from the other neighbouring genes. This approach has proven to be highly effective for stabilizing dispersion estimates when the sample size is small. Initially, a preset weighting was used for the prior distribution [7]. Later, a QL strategy using the deviances was used to optimize the prior weight, leveraging limma’s hyperparameter estimation algorithm [9]. A variation of the weighted likelihood empirical Bayes algorithm that is more robust to observation outliers was also implemented [31].

#### Quasi-likelihood

edgeR v3 introduced the QL generalization of the NB model [19, 10]. In edgeR’s QL pipeline, the NB-dispersions are used to model the global trend in biological variation while the quasi-dispersions are used to accommodate gene-specific variability. In edgeR v3, the trended NB-dispersions from the weighted likelihood approach described above were generally used. In edgeR v4, a constant NB-dispersion estimated from the most highly expressed genes is used by default. In either case, the quasi-dispersions are then estimated by the same empirical Bayes moderation strategy as in limma but with the GLM residual deviances in place of the residual variances used by limma [23]. As in limma, the posterior quasi-dispersion estimates are moderated towards a trended prior with prior degrees of freedom (df) estimated from the data as part of the empirical Bayes procedure. The empirical Bayes procedure optionally includes robust hyperparameter estimation, which protects against hypervariable genes [23], and the ability to model the quasi-dispersion trend in terms of a prior precision estimator.

The precision of the posterior quasi-dispersion estimators is summarized by their posterior df, which is made up of the prior df plus the residual df. Quasi-F-tests are then used to test hypotheses with the posterior df entering as the residual df. The quasi-F-tests provide more rigorous error rate control than other analysis methods that do not fully reflect the uncertainty with which the quasi-dispersions are estimated [19, 41]. QL is now the default recommended pipeline in edgeR because of its robustness and very reliable FDR control [10, 24, 25].

#### Hypothesis testing for general contrasts

edgeR v1 was able to make pairwise comparisons between treatment groups, but later versions of edgeR can test very general null hypotheses specified by any linear contrast of the linear model coefficients. This provides enormous flexibility for users. For example, one might compare one group to the average of other groups simply by specifying suitable contrast weights for the means of the different groups. Tests can be conducted for a single contrast or for several contrasts at a time, leading to an analysis of deviance test on several df [8, 10]. For example, one can conduct an ANOVA-like test for differences between several groups by specifying any set of contrasts that distinguish the groups. The edgeR GLM functions *glmFit* and *glmLRT* conduct likelihood ratio tests [7]. The edgeR QL functions *glmQLFit* and *glmQLFTest* conduct quasi F-tests where the likelihood ratio statistic is the numerator of the F-statistic and the quasi-dispersion is the denominator [8, 10].

#### Supporting fractional counts

edgeR was designed for analysing digital gene expression data, which usually comes in the format of integer counts, and the NB distribution has a discrete probability mass function defined on the non-negative integers. Tools that count reads overlapping genomic features such as featureCounts [42] or HTSeq [43] do indeed produce integer counts. However, the read counts can be fractional when the number of sequence reads originating from a gene or transcript is estimated in a probabilistic manner. RSEM, kallisto and Salmon are examples of software tools that output estimated gene and transcript counts that are not integers [44, 45, 46].

To handle fractional data without the need for rounding, a continuous generalization of the NB distribution has been implemented in edgeR. All the binomial coefficients of the NB probability mass function were replaced with gamma functions, resulting in a probability function that is equal to NB probabilities when evaluated at integers but which varies smoothly and continuously for intermediate values. This generalization ensures that the results returned by edgeR are unchanged for integer counts but vary smoothly and gradually for fractional counts.

#### Correcting quasi-likelihood for bias

Traditional quasi-likelihood statistical theory relies on a chisquare approximation to the GLM deviances, which can be justified by a saddlepoint approximation [40, 8]. The chisquare approximation to NB deviances is excellent when the NB-dispersion is small and the counts are large. Unfortunately, for single-cell data especially, the NB-dispersions are often large and the counts may be small. For genes with average count below 1, the chisquare approximation substantially underestimates the quasi-dispersions.

In edgeR v3, we already implemented an adjustment for the residual df when some of the fitted values were exactly zero [21]. In edgeR v4, we have implemented a more comprehensive approach. We drop the use of the saddlepoint approximation and instead approximate the unit deviances by scaled chisquare random variables on fractional df. The df and the scaling factor are chosen to match the first two moments (the mean and variance) of the unit deviances exactly, given the fitted mean and the NB-dispersion for each observation. This approach yields very nearly unbiased quasi-dispersion estimators even for fitted values close to zero and for quite large NB-dispersions (Figure 1). The new approach has been shown to improve performance for transcript differential expression in particular [25].

**Figure 1:**
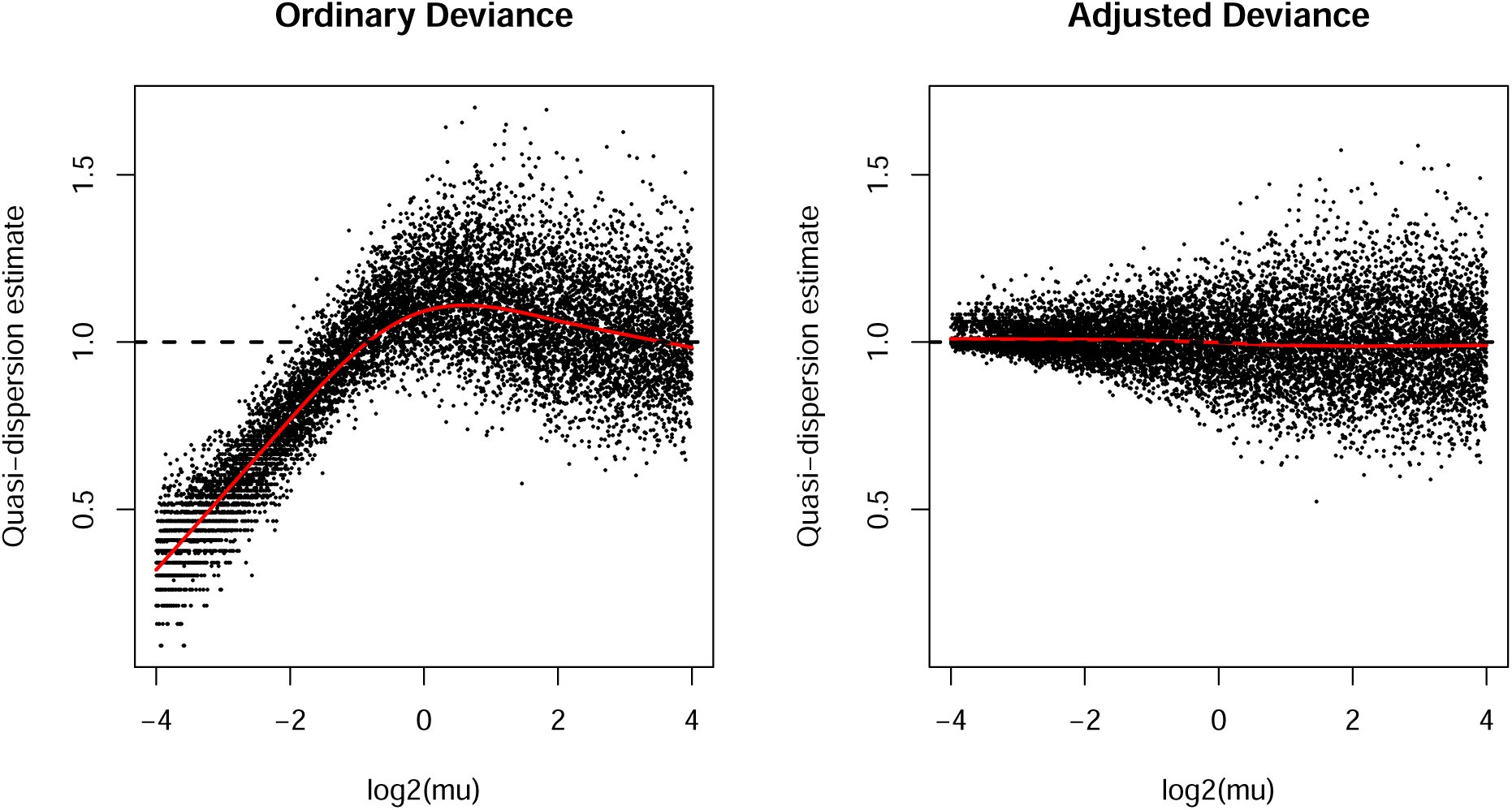
Bias-adjusted deviances produce unbiased quasi-dispersions. Poisson counts were generated for 100 samples and 10,000 genes. The true quasi-dispersion is 1 for all genes. The left panel shows quasi-dispersions estimated using the mean-deviance estimator from classical generalized linear model theory. The right panel shows the mean-deviance estimators for the same data using the bias-adjusted deviances from edgeR v4. The ordinary deviances have negative or positive biases, depending on the expected count size, whereas the edgeR v4 adjusted deviances are unbiased for all count sizes.

The bias-corrected deviances produce residual df that are fractional and, in some cases, can be less than 1. In order to support edgeR v4, limma’s empirical Bayes hyperparameter estimation was revised in limma 3.62.0 to give improved performance when the residual df are unequal between genes and possibly small. The new methodology has been implemented in limma’s *fitFDistUnequalDF1* function, which is called automatically by *glmQLFit* in edgeR v4.4.0 as part of the new QL pipeline.

#### Larger datasets

The edgeR v4 QL pipeline also allows edgeR to analyse larger datasets than previously. The new QL pipeline allows the *estimateDisp* function to be bypassed resulting in substantially reduced computation time for large datasets [25]. A oneway multi-group analysis with 1000 samples and 10000 genes takes about 30 seconds on a laptop computer including dispersion estimation, GLM fitting and testing for differential expression. The same data with a more complex design matrix takes less than two minutes, depending on the number of predictors.

### Analysing next generation sequencing data

#### Standard analysis steps

edgeR offers a complete differential analysis pipeline from read counts to lists of differential genes and pathways. The standard steps in most analyses include data import, filtering out lowly expressed genes, normalization, data exploration, dispersion estimation, fitting GLMs and hypothesis testing. edgeR implements one or more functions for each of these steps (Figure 2).

**Figure 2:**
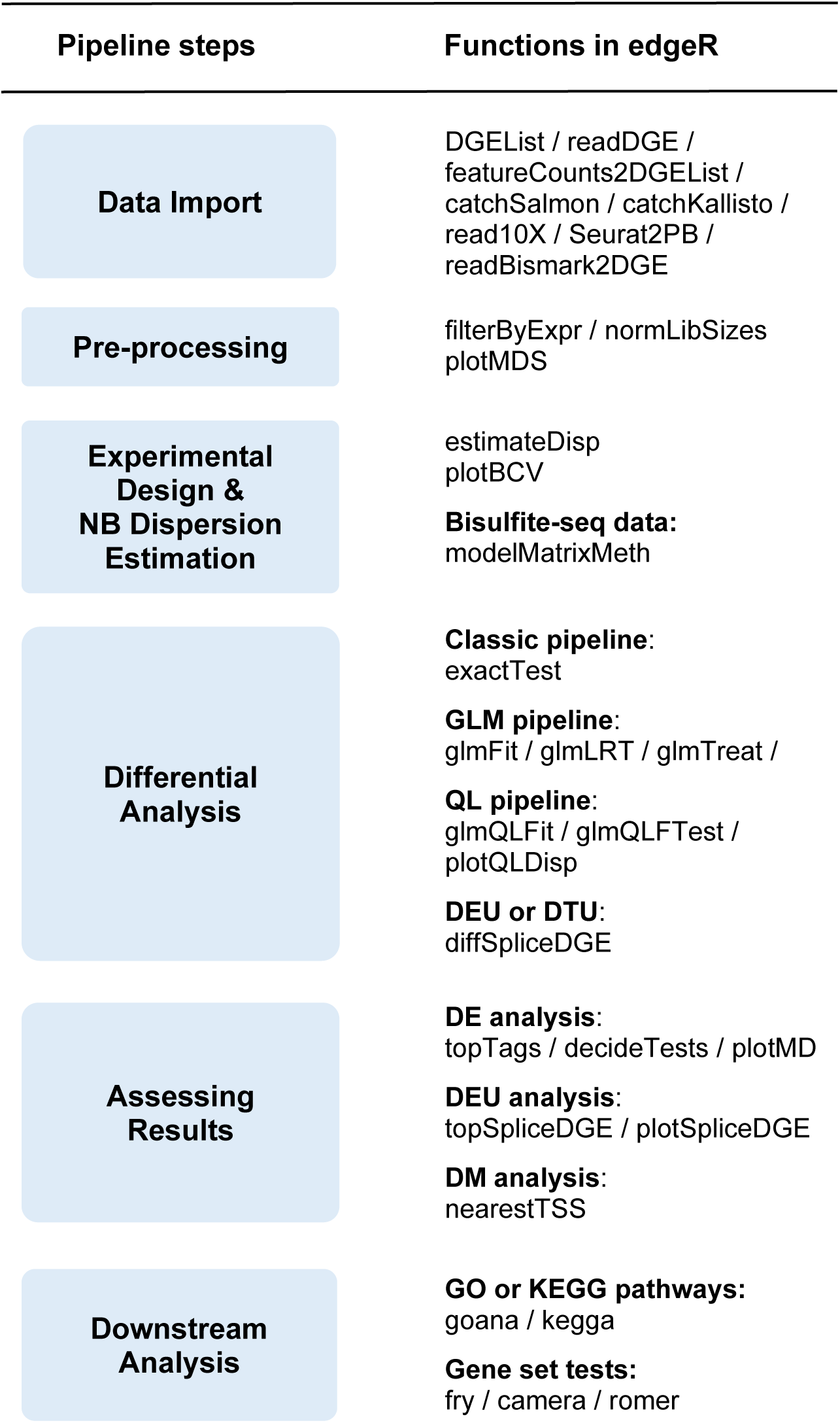
The edgeR workflow. The diagram shows the main steps in an edgeR analysis. Individual functions involved in each step are shown on the right.

#### Data import

The minimum data that edgeR needs to conduct an analysis is a numeric matrix of counts and a factor or covariate distinguishing the samples. For most users, the first step is to assemble the read counts and annotation into a ‘DGEList’ object, which is edgeR’s data container. If users already have a numeric matrix in R, then the *DGEList* function converts it to a ‘DGEList’. If users have imported the counts and gene annotation from a text file into an data-frame using base R, then *DGEList* will process the data-frame and attempt to distinguish the counts from the annotation columns. edgeR’s readDGE function will collate read count files written by Subread-featureCounts [42] or by HTSeq-count [43], both of which write one file per sample. Alternatively, the *featureCounts2DGEList* function imports output from Rsubread’s *featureCounts* function [47] directly without the need to write intermediate files to disk.

10x Genomics Chromium is the most popular platform for single-cell RNA-seq. edgeR’s *read10X* function reads output from the 10x Genomics Cellranger pipeline [48] and assembles the UMI counts into an integer matrix in R with accompanying cellular barcodes and gene symbols. Seurat is an extremely popular R package for statistical analysis of scRNA-seq [49]. edgeR’s *Seurat2PB* function imports data from Seurat and aggregates it to pseudo-bulk form suitable for an edgeR analysis [50]. RSEM [44], kallisto [45], and Salmon [46] have become popular for quantifying RNA-seq data because of their ability to estimate transcript-level expression. edgeR’s *catchRSEM*, *catchKallisto* and *catchSalmon* functions import output from these tools, read in transcript counts and estimate the assignment uncertainty for each transcript.

Bismark is a popular software tool for read mapping and methylation calling of bisulfite sequencing (BS-seq) data [51]. edgeR’s *readBismark2DGE* function reads Bismark coverage files and collates the methylated and unmethylated read counts from multiple files into a ‘DGEList’ object [13].

#### Filtering low count features

Genes with very low read counts are filtered before downstream analysis. Filtering is done partly because very low expressed genes are of little biological interest and because NB-dispersion estimation is unreliable for such genes. The main reason however is that genes with too few reads cannot achieve statistical significance even if they truly are differentially abundant. Keeping such genes in the downstream analysis increases the amount of multiple testing and decreases statistical power without any compensating advantage. edgeR’s *filterByExpr* function is used to keep only those genes or features that have sufficient counts to achieve statistical significance when meaningful differential abundance is present.

#### Effective library sizes and the offset matrix

edgeR’s *normLibSizes* function estimates an effective library size (ELS) for each sample. It estimates a normalization factor for each sample, which multiplies the raw library size to become the ELS. TMM (trimmed mean of M-values) [52] is the default normalization method but a new TMMwsp method (TMM with singleton pairing) has been added to provide more robust behaviour for sparse data with many zeros. The TMM method discards genes with zero count in either library when comparing each library to the reference library. TMMwsp rescues genes with a positive count in just one of the two libraries (a singleton) by pairing them up with another gene with a singleton count in the other library.

The log-ELSs become GLM offsets in downstream edgeR analyses. The offset matrix can also be set directly, which provides a way to apply nonlinear normalization procedures or to reflect biases at the observation level that arise from factors such as GC content or gene length. The *normalizeBetweenArrays* method for ‘DGEList’ objects implements quantile or cyclic loess normalization in this way, methods traditionally associated with microarrays. edgeR supports custom normalization procedures implemented by external packages such as cqn [28], EDAseq [27] or tximport [30], by importing the appropriate offset matrix. edgeR’s *scaleOffset* function ensures that the scale of externally defined offset matrices are consistent with library sizes while preserving the normalization results from those external methods.

#### Model fitting and differential analysis

edgeR defines linear models in the same way as limma [53]. Users create a design matrix appropriate for their experimental design using standard base R functions. edgeR has the flexibility to work with any full-rank design matrix with any number of covariates or factor effects.

In edgeR v4, one can proceed directly from normalization to *glmQLFit* to fit GLMs and *glmQLFTest* to test hypotheses. In edgeR v4, the *glmQLFit* function estimates both NB and quasi-dispersions automatically.

In earlier edgeR versions, the NB-dispersions are first estimated by *estimateDisp* [9]. Then one proceeds to *exactTest* in the classic pipeline, to *glmFit* followed by *glmLRT* in the GLM pipeline, or to *glmQLFit* followed by *glmQLFTest* in the QL pipeline.

The functions *glmFit* and *glmQLFit* shrink the estimated log-fold-changes slightly towards zero to avoid infinite values and to improve prediction accuracy. edgeR implements the concept of a “prior count” to quantify the prior information determining the amount of shrinkage [54].

For all pipelines, the number of differential genes at any given FDR threshold can be shown by the *decideTests* function and the top differential genes can be displayed by the *topTags* function.

#### Testing differences relative to a fold-change threshold

The *glmLRT* or *glmQLFTest* functions can be replaced by *glmTreat* if one wants to test differential abundance relative to a higher fold-change threshold.

The standard edgeR analysis identifies differential expression based on statistical significance regardless of how small the difference might be. There are circumstances where researchers are interested in studying genomic features of which the expression levels change by a certain amount. A few *ad hoc* approaches have been used to select features with large fold-changes. For example, some studies applied a fold-change cut-off and then ranked all the genes above that fold-change threshold by p-value. There were also some cases where genes were first chosen according to a p-value cut-off and then sorted by their fold-changes. However, these *ad hoc* approaches tend to prioritise low expression genes and can lead to loss of FDR control.

To assess differential expression relative to a threshold in a statistically rigorous way, the TREAT method was developed under the limma empirical Bayes framework [55, 23]. It adopts a re-centred moderated t-statistic to provide an upper bound of the type I error rate. This approach yields an easily computable conservative p-value for testing against a fold-change threshold. We adopted the idea of TREAT and extended it to the edgeR NB GLM framework. Unlike TREAT, the edgeR approach computes the expectation rather than the maximum value of the type I error rate as the p-value of the test, a refinement that increases the statistical power of the test while still controlling the FDR. This method is implemented in the *glmTreat* function in edgeR, and it can be used under both the likelihood ratio test (LRT) and the QL F-test pipelines.

#### Gene set enrichment analysis

It is often helpful to interpret differential results in terms of higher-order biological processes. The limma package [17] provides a number of functions to facilitate this interpretation in terms of standardized gene annotation or gene sets. The *goana* and *kegga* functions determine the overlap of a list of differential genes with categories in the Gene Ontology (GO) database [56] or with pathways in the Kyoto Encyclopedia of Genes and Genomes (KEGG) [57] database. The *goana* and *kegga* functions evaluate the significance of the overlaps with hypergeometric tests, optionally adjusting for any power trend bias associated with average expression or with gene length [58]. Alternatively, differential analyses can be conducted directly for pre-defined gene sets such as the Molecular Signatures Database collections [59] or gene sets from previous independent studies. The *roast*, *mroast* and *fry* functions perform self-contained tests to assess whether the majority of the genes in a set are differentially expressed across the comparison of interest [60]. The *camera* function performs a competitive gene set test to find gene sets that are highly ranked relative to other genes in terms of differential expression [61]. The *romer* function conducts a gene set enrichment analysis analogous to the GSEA approach [62]. The functions all achieve rigorous FDR control even in the presence of inter-gene correlations [60, 61].

edgeR v4 includes S3 methods for all the above gene set functions. The *goana* and *kegga* methods operate on an ‘DGEExact’ or ‘DGELRT’ object produced by *exactTest*, *glmLRT* or *glmQLFTest*. They automatically extract differentially expressed genes under a given significance threshold and perform generalized hypergeometric tests for enrichment of GO terms or KEGG pathways in the list of differentially expressed genes.

edgeR’s *roast*, *mroast*, *fry*, *camera* and *romer* methods operate on ‘DGEList’ data objects and use a z-score strategy to translate negative binomial counts into normal deviates suitable for limma analyses. The functions fit NB GLMs using the null hypothesis design matrix to the count data and convert the counts to their z-score equivalents using a quantile-quantile transformation with the estimated means and dispersions. edgeR supports integer or non-integer data in computing the z-score equivalents. For integer values, a continuity correction is applied by splitting the probability mass of each integer in two. It computes the mid-p tail probability of the given quantile and then converts it to the standard normal deviate with the same cumulative probability distribution value [63]. Non-integer values are handled by interpolation. In the edgeR context, the *fry* function is always preferred over *roast* or *mroast*. In the limma context, *fry* implements an analytic approximation to *mroast* that is faster and yields higher resolution p-values. In edgeR, the z-score transformation ensures that the distributional assumptions of *fry* are satisfied exactly.

#### Transcript-level differential expression

edgeR versions v1–v3 were designed for gene-level analyses of RNA-seq, but edgeR v4 adds the ability to conduct differential expression analyses for individual transcripts (isoforms) of each gene using output from RSEM [44], Salmon [46] or kallisto [45]. Transcript quantifications are inherently more uncertain than gene-level read counts because of ambiguous assignment of RNA fragments to isoforms [30]. Whereas sequence reads can usually be assigned unambiguously to a gene, reads are very often compatible with multiple transcripts for that gene, particularly for genes with many isoforms. This read to transcript ambiguity (RTA) causes technical overdispersion in the read counts and the amount of overdispersion depends on the degree of overlap of each transcript with other transcripts rather than on its expression. The edgeR gene-level differential expression pipeline does not perform optimally on transcript counts because the RTA disrupts the mean-variance relationship normally observed for gene level RNA-seq data and therefore interferes with the efficiency of the empirical Bayes dispersion estimation procedures. However, the RTA dispersion can be modelled elegantly using overdispersed Poisson distributions in accordance with edgeR’s QL model.

Salmon and kallisto have the ability to repeatedly resample each RNA-seq library to generate bootstrap samples, and this resampling provides a means to estimate technical variability. Alternatively, RSEM and Salmon can generate Gibbs samples from the posterior distribution, which can be used in the same way as bootstrap samples. The edgeR functions *catchRSEM*, *catchKallisto* and *catchSalmon* read the outputs and use the bootstrap or Gibbs samples to estimate a quasi-dispersion arising from RTA for each transcript. The quasi-dispersions can then be divided out of the transcript counts, as described in Methods, leading to divided counts that can be analysed by edgeR’s gene-level software tools with full statistical efficiency [24, 25]. The quasi-dispersions estimate the variance-inflation induced by RTA and scale down the transcript counts so that the resulting library sizes reflect their true precision. The divided counts follow the traditional NB mean-variance relationship so that standard methods designed for gene-level differential expression analyses can be applied without further modification. The divided counts are not integers but are handled by edgeR’s continuous generalization of the NB distribution.

The divided transcript counts can also be input to edgeR’s *diffSpliceDGE* function for a differential transcript usage analysis, as described in the next section.

#### Differential exon usage

RNA-seq is a powerful tool for studying alternative splicing, whereby exons are differentially combined or skipped, resulting in multiple isoforms encoded by a single gene. One way to detect alternative splicing events is to test for differential exon usage (DEU), which allows fast detection of potentially differentially spliced genes without identifying the actual isoforms present in different groups.

In edgeR, DEU analysis can be performed using the *diffSpliceDGE* function. The function accepts a matrix of exon read counts and compares the change of the expression level of each exon to the change of the expression level of the gene containing that exon under a certain comparison. NB GLMs are fitted to the exon counts and exon-level NB-dispersions are estimated, as would be done for a gene-level analysis but at the exon level.

Depending on the pipeline of choice, statistical tests can be performed using either likelihood ratio tests or quasi-likelihood F-tests to obtain a test statistic and a p-value for each exon. The *diffSpliceDGE* function offers two different ways to provide inferences at the gene level. The first approach is to combine the exon-level test statistics for all the exons within a gene and then to conduct a gene-level test. The second approach is to combine the exon-level p-values using Simes’ method for all the exons within a gene to get a p-value for that gene [64]. The first method favours genes for which many exons are differentially spliced. The Simes’ method, on the other hand, is likely to be more powerful for picking up genes for which only a minority of the exons are differentially spliced.

#### shRNA-seq screens

edgeR can be used to analyse data from CRISPR-Cas9 and shRNA-seq genetic screens to identify the change of single guide RNA (sgRNA) or short hairpin RNA (shRNA) in the selected cell population relative to a control population [65]. Given a list of sample index sequences and sgRNA- or shRNA-specific sequences from an amplicon sequencing screen, the edgeR *processAmplicons* function obtains read counts for each sgRNA/shRNA in the screen across all samples. This is done by counting the number of times each sample index and sgRNA/shRNA combination could be matched in reads from the input fastq files. These read counts are then organized and stored in a ‘DGEList’ object for a standard edgeR downstream DE analysis, which identifies a list of sgRNAs/shRNAs that are differentially expressed between the groups. To interpret the results at gene level, multiple sgRNAs/shRNAs that target the same gene are grouped together and treated as a ‘set’. Gene set testing methods such as *camera* and *fry* can then be applied to summarize the results at gene level [60, 61].

#### Differential methylation analysis

Bisulfite sequencing (BS-seq) has become the gold-standard technology for studying DNA methylation [66]. A number of software tools, such as Bismark, have been developed to align BS-seq reads to a reference genome and count the number of C-to-T conversions at each CpG locus in each sample [51]. Downstream analyses often focus on the identification of CpG loci or regions that are differentially methylated between different experimental conditions.

Since the output count matrix from methylation calling software contains both methylated and unmethylated Cytosines at each CpG locus, the structure of BS-seq data is analogous to paired-samples RNA-seq data. This allows the differential methylation analysis to be conducted using existing edgeR pipelines developed for RNA-seq differential expression analyses [13]. In particular, edgeR models both methylated and unmethylated counts as NB distributed, and the variation between replicate samples is captured by the NB-dispersion parameter. One key difference between BS-seq and paired-samples RNA-seq analysis is that the pair of libraries that hold the methylated and unmethylated read counts from each sample are treated as a unit, and hence share the same library size. A special design matrix constructed by the *modelMatrixMeth* function includes individual sample effects accounting for read coverage plus group-specific coefficients representing the log-ratio of methylated to unmethylated reads for each group. The subsequent analysis is identical to any other edgeR analysis, thus giving methylation analysts access to the full downstream capabilities of the edgeR package. The edgeR methylation approach was originally developed for reduced representation BS-seq, but has been extended to whole-genome BS-seq and shown to outperform competing methods [67].

Likelihood ratio or quasi-F tests can be performed to identify CpG loci at which the proportion of methylated reads are significantly different between the groups. Given the fact that unmethylated CpGs are often enriched in gene promoters [68, 69, 70, 13], a gene oriented differential methylation analysis can be conducted by aggregating the methylated and unmethylated CpG counts in gene promoter regions [13]. This improves the statistical power of differential methylation analysis and provides an interpretation at gene level.

The same edgeR analysis strategy as for BS-seq can be applied to any type of sequencing application where the sequence reads are classified into two classes at each genomic loci, and the aim is to test for changes in the relative proportions of the two classes. The authors have used this approach to test for changes in haplotype-specific expression, for example.

#### Single-cell pseudo-bulk analysis

Single-cell technology has revolutionized biomedical research, and many R packages and software tools have been developed for analysing scRNA-seq data in the past few years [71, 72, 49]. As the technology evolves and costs reduce, single cell experiments with biological replicates have become more standard.

Accounting for biological variation between replicates is crucial for single-cell differential expression analyses so that the results are not driven by particular samples. One popular approach to do so is the pseudo-bulk method, whereby read counts of the cells within the same cluster and from the same sample are aggregated together to form pseudo-bulk samples [50]. The edgeR differential expression analysis pipeline can be applied to pseudo-bulk samples for identifying marker genes of each cell cluster.

Pseudo-bulk analysis by edgeR has been shown to give better performance than other methods for single-cell differential expression analysis in the presence of replicate samples [50]. This approach was applied to a recent large cohort single-cell study, and successfully identified marker genes of different cell populations while accounting for the biological variation between tissue specimens [73].

### Graphic exploration

#### MDS plot

edgeR offers diagnostic plots for data exploration (Figure 3). One of the most commonly used plots is the multi-dimensional scaling (MDS) plot, generated by *plotMDS*, which visualizes the similarities and dissimilarities between samples on a two-dimensional scatter plot (Figure 3a). Distances on the MDS plot represent leading log2-fold-changes, defined as the root-mean-square average of the top largest log2-fold-changes, between each pair of samples. The percentage variation explained is also calculated and returned for the selected dimensions. MDS plots are an unsupervised visualization method that provides a easy and fast way to check the relationship of all the samples in a data set. limma’s *removeBatchEffect* can be used in conjunction with an MDS plot to display treatment or group effects after adjusting for batch effects or covariates.

**Figure 3:**
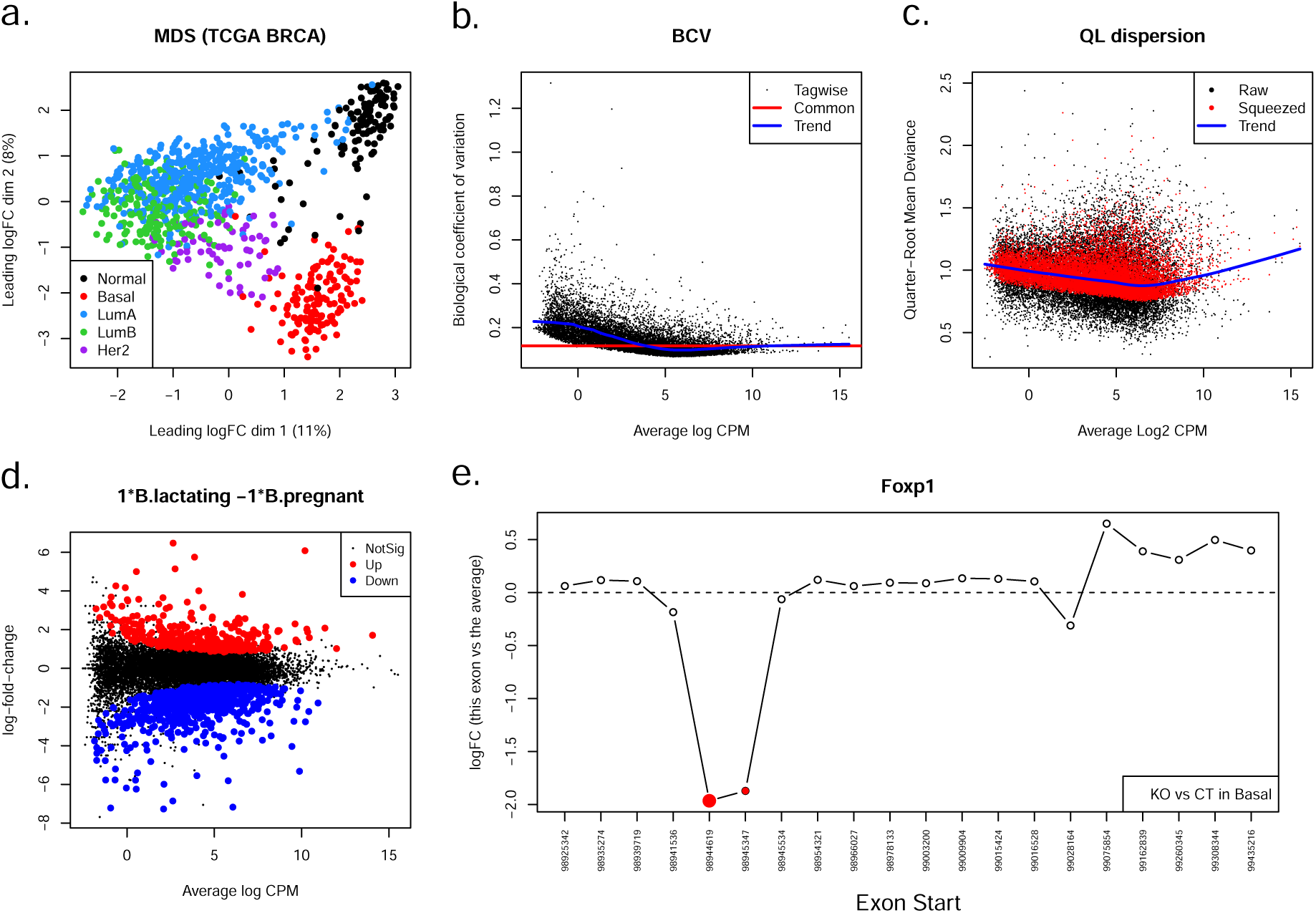
Example diagnostic plots produced by edgeR. **(a)** Multidimensional scaling (MDS) plot of the human TCGA breast cancer data set of 882 individual RNA-seq samples, generated by the *plotMDS* function. Samples are coloured by breast cancer subtype identified by PAM50 signatures. Samples of the same cancer subtype are clustered together on the MDS plot. The proportions of variance explained by the first two dimensions are also shown. **(b)** Scatter plot of the biological coefficient of variation (BCV) against the average abundance of each gene in the mouse mammary gland RNA-seq data set, generated by the *plotBCV* function. The plot shows the square-root estimates of the common, trended and tagwise NB-dispersions. **(c)** Scatter plot of the quarter-root quasi-dispersion against the average abundance of each gene in the mouse mammary gland RNA-seq data set, generated by the *plotQLDisp* function. The “Raw” and “Squeezed” values represent the dispersion estimates before and after the empirical Bayes moderation towards the trend. **(d)** MD plot showing the log-fold change and average abundance of each gene between the lactating and pregnant samples in the basal cell population, generated by the *plotMD* function. Significantly up and down DE genes are highlighted in red and blue, respectively. **(e)** Plot showing relative log-fold changes by exons for the *Foxp1* gene in an RNA-seq experiment where two of the exons were silenced in the KO group. The plot is generated by the *plotSpliceDGE* function. The relative logFC is the difference between the exon’s logFC and the overall logFC for the gene, as computed by the *diffSpliceDGE* function. The significant differentially used exons are highlighted in red. The size of the red dots are weighted by its significance. The start position of each exon is labelled at the bottom.

#### Dispersion plots

In a standard analysis pipeline, edgeR provides three different types of NB-dispersion estimate for each gene [7]. The first is the common dispersion, the global dispersion estimate across all genes. The second is the trended dispersion where the dispersion is predicted from the abundance of each gene. The third is the gene-specific dispersion for each individual gene estimated by weighted likelihood empirical Bayes. These dispersion estimates can be visualized in a BCV plot generated by the *plotBCV* function in edgeR (Figure 3b). The y-axis of the BCV plot displays square-root NB-dispersion, which can be interpreted as biological coefficient of variation (BCV).

Under the QL framework, gene-specific variability is measured by the quasi-dispersion. The quasi-dispersion estimates can be visualized in a plot generated by the *plotQLDisp* function (Figure 3c). The plot displays the global QL mean-dispersion trend, the raw quasi-dispersion estimates, and the squeezed estimates after empirical Bayes moderation.

#### MD plots

Mean-difference plots (MD plot) are useful for assessing the differences between samples or groups [17]. To examine each individual sample in a data set, an MD plot can be constructed by comparing the log-expression values of all the genes in that sample with the mean of those in all other samples. The skewness of the log-ratios on an MD plot would suggest that a normalization is required to correct for the compositional biases. MD plots can also be useful tools to spot biases that are typically observed in DNA-based sequencing experiments, such as immunoprecipitation efficiency biases and also more complex trended biases for which the magnitude of the systematic differences between samples will change with the average abundance [11]. When a differential expression analysis is performed, an MD plot can be generated to show the relationship between the log fold changes and the average log-CPM values for all the genes under that comparison (Figure 3d). This allows a quick identification of outlier genes that change substantially under the comparison. MD plots are implemented in limma and edgeR by *plotMD*, a generic function with methods for edgeR data objects and fitted model objects.

#### Splicing plot

Results from *diffSpliceDGE* can be visualized in a differential splicing plot generated by the *plotSpliceDGE* function (Figure 3e). In a splicing plot, relative log-fold changes by exons for the specified gene are plotted for all the exons within the gene from left to right in the order of their genomic locations. Exons that are significantly differentially used are highlighted in red. This easy and simple visualization approach provides a quick way to examine the differences of gene structure between the groups.

### Computational efficiency

#### C implementation

edgeR typically fits NB GLMs to tens of thousands of genes in a given dataset. For some applications, the number of genomic features may in the hundreds of thousands, even millions are possible. The vector of GLM parameters needs to be estimated for each individual gene using an iterative computational algorithm. The iteration must reliably converge and provide accurate estimates for every gene. When the NB-dispersions are estimated, the genewise GLMs need to be refitted repeatedly with different candidate values for the dispersions. The efficiency and reliability of the iterative GLM fitting is therefore a core factor in determining the overall computational load of the package.

The classical algorithm used for GLM fitting is an iteratively reweighted least squares iteration that is equivalent to Fisher-scoring [8]. The basic algorithm is adequate for classical univariate applications but is not sufficiently reliable in the edgeR context where even a small percentage of convergence failures would be problematic.

The original GLM version of edgeR v2 was written purely in R. If the experiment design could be transformed to a oneway layout, then unmodified Fisher scoring was used which, in this scenario, is fast and reliable. For other designs, a novel GLM algorithm was implemented using a simplified approximation to the Fisher information matrix and a line search strategy to ensure convergence [7]. To achieve acceptable computational speed, the R implementation was vectorized so that the NB GLMs were fitted for all the genes simultaneously.

Starting in edgeR v3, the low-level GLM fitting functions were re-implemented in C and C++, with dependence on the Rcpp package to simplify memory management [74]. The simplified Fisher-scoring with line search was replaced by full Fisher-scoring with a Levenberg-Marquardt modification to ensure convergence [75, 76]. The direct C/C++ implementation increases speed and allows observational weights to be supported. The improved speed and numerical stability of the C code allows more Fisher-scoring iterations to be performed, leading to more accurate final estimates. Other edgeR components rewritten in C/C++ included the Cox-Reid APL, the GLM deviances, loess curves, and maximizing the interpolant as part of the weighted likelihood empirical Bayes. In edgeR v4.4.0, the code has been converted to pure C, in the same style as for base packages in R, and dependence on Rcpp is no longer required.

#### CompressedMatrix class saves memory

When fitting the genewise GLMs, the offsets log *L_gi_*, weights *w_gi_* and NB-dispersions *ϕ_g_* all enter into the calculations and are treated as known values. The offsets are often the same for every gene but can be observation-specific. The weights are usually constant (*w_gi_* = 1) but could be sample-specific or observation-specific. The NB-dispersions are typically gene-specific but might be constant and could in principle be observation-specific. In other words, each of these quantities could be a constant, a row vector, a column vector, or a matrix. It is convenient to represent each of these quantities as a gene×sample matrix in R, but doing so is memory inefficient if the same numerical values are repeated over genes or samples or both.

In edgeR v4, a simple but elegant object class called ‘CompressedMatrix’ has been introduced to make these matrices more efficient. A ‘CompressMatrix’ object is subsettable in R as a matrix but, internally, stores only the minimum information required to construct the matrix from the repeat structure. The minimum information can be a constant, a row vector, a column vector, or a complete matrix. This simple but efficient representation saves considerable memory for most analyses when the offsets, weights and NB-dispersion are passed to C or stored in the fitted model ‘DGEGLM’ object.

### User interface

#### Object-oriented programming

An edgeR analysis consists of a number of distinct steps (Figure 2). Each of the major steps in the pipeline — (1) data import, (2) model fitting, (3) statistical testing and (4) gene ranking — produces a classed R object that can be input to functions downstream in the pipeline.

Data import produces an object of class ‘DGEList’, which stores the read counts and associated information (Figure 4a). The essential components of a ‘DGEList’ object are the matrix of raw counts and a data-frame containing the sample information. Other optional components include gene annotation and a design matrix.

**Figure 4:**
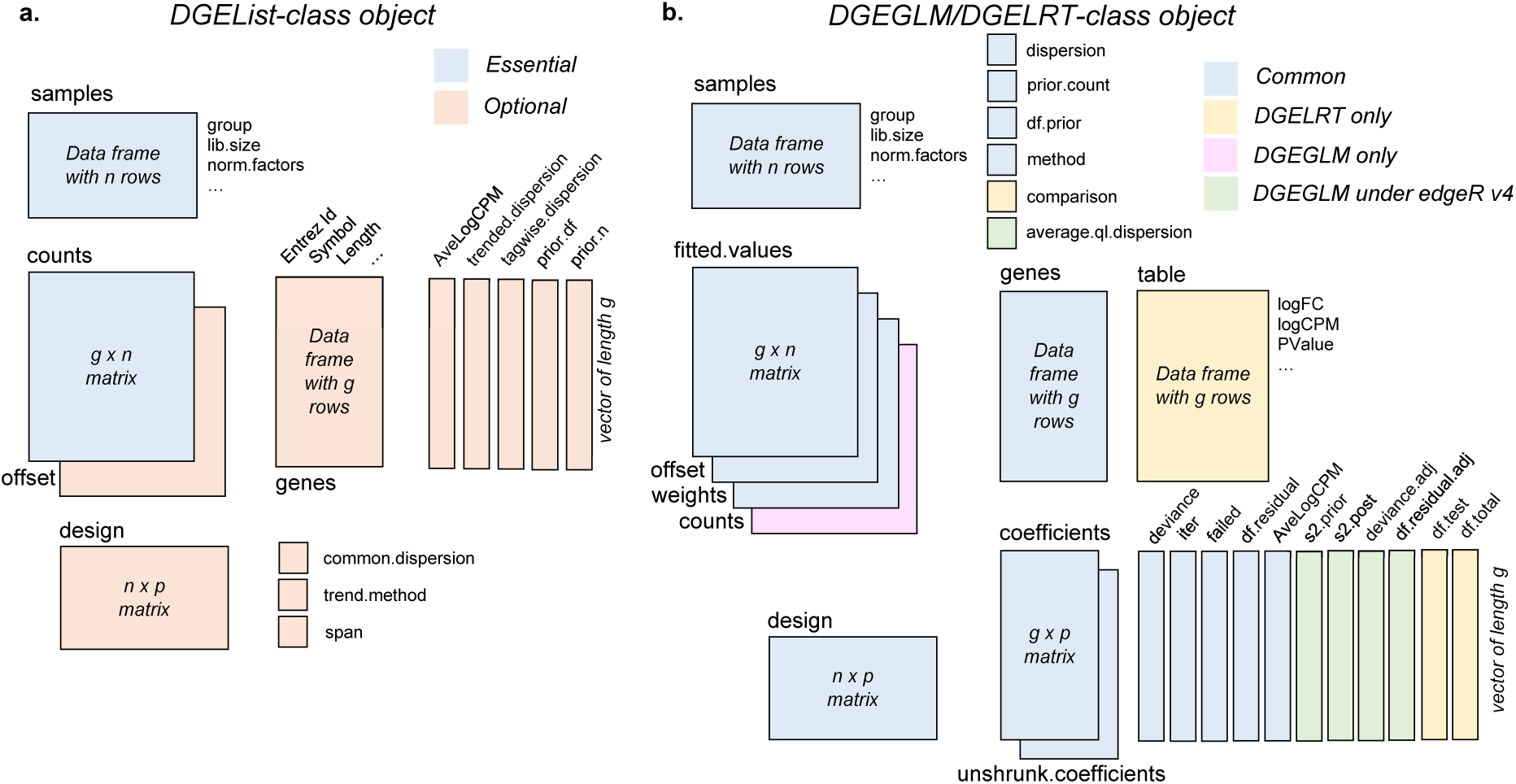
The data structure of **(a)** DGEList-class object, and **(b)** DGEGLM-class and DGELRT-class object.

Normalization adds library-size normalization factors or offsets to the ‘DGEList’ object and dispersion estimation adds NB-dispersion estimates.

Model fitting by *glmFit* or *glmQLFit* produces an object of class ‘DGEGLM’, which stores the generalized linear model fits and dispersion estimates (Figure 4b). The ‘DGEGLM’ object includes GLM deviances, coefficient estimates, and fitted values, plus all the parameters used in the fitting process. The object produced by *glmQLFit* contains all the same information as *glmFit* plus additional components storing the quasi-dispersion estimation.

Testing for differential abundance produces either a ‘DGEExact’ object, if exact NB tests were used, or a ‘DGELRT’ object if GLMs were used (Figure 4b). The object includes a data-frame table containing the genewise log-fold changes, average log-CPMs, test statistics and p-values for the comparison specified. The ‘DGELRT’ output object preserves most of the components of the input ‘DGEGLM’ object but drops the read counts if they were present.

Finally, a ‘TopTags’ object containing a ranked gene list is created by the *topTags* function. The ‘TopTags’ object contains a sorted data-frame of genewise tests results including adjusted p-values that adjust for multiple testing. The default adjustment method controls the FDRs for a ranked list using the Benjamini and Hochberg method [77].

All of these data classes obey many analogies with matrices. For ‘DGEList’, rows correspond to genes/features and columns to different samples. For ‘DGEGLM’, rows correspond to genes/features and columns correspond to linear model coefficients. For ‘DGEExact’, ‘DGELRT’ and ‘TopTags’, the columns correspond to the differential results table. The standard R functions *summary*, *dim*, *length*, *ncol*, *nrow*, *dimnames*, *rownames*, *colnames* have methods for each of these classes. ‘DGEList’ objects can be subsetted by rows and columns as for a matrix. Multiple ‘DGEList’ objects can be row-binded or column-binded together, again as for matrices. The other edgeR classes can be subsetted by rows, but column subsetting is disallowed because it would not produce a valid object of that class.

All edgeR objects can be coerced to a data-frame using *as.data.frame* in R. All edgeR classes except ‘TopTags’ belong to the virtual class ‘LargeDataObject’ for which a show method is defined to display the leading rows of each component vector, matrix or data.frame.

A design aim of edgeR is that users should be able to easily explore and manipulate all objects produced by edgeR functions and that doing so should require a knowledge of base R only. While all the edgeR objects are formally registered using R’s S4 system, the objects are also fully manipulatable as ordinary R lists and the list components are standard base R objects: data-frames, vectors and matrices. This is analogous to the behaviour of core R modelling functions such as *lm* or *glm*. At each stage of the edgeR pipeline, functions have S3 object-orientated methods that accept objects from upstream in the pipeline, but also have default methods that accept input in the form of atomic R objects. This allows direct programming access to any stage of the edgeR pipeline from external software tools.

In edgeR v4, support has been added for the popular ‘SummarizedExperiment’ object class developed by the Bioconductor core team (https://bioconductor.org/packages/-SummarizedExperiment) [18]. All relevant edgeR functions now accept raw data in a ‘SummarizedExperiment’ container that includes a ‘counts’ assay, making it simpler to access edgeR’s functionality in the context of other Bioconductor pipelines using ‘SummarizedExperiment’.

#### Documentation

The edgeR package is supported by extensive documentation, examples and online help. edgeR v4.4.0 documents 253 exported functions or methods, each of which has a detailed documentation help page including example code and information on distinct methods for generic functions. Every help page formally documents the class of each input argument and the class and structure of the output object. edgeR v4.4.0 includes a 138-page User’s Guide. It covers a wide range of topics such as the underlying statistical framework of edgeR, normalization, different analysis pipelines, setting up appropriate design matrices and downstream analysis. The edgeR User’s Guide also provides 10 individual case studies with complete data source and analysis R code. These fully worked case studies cover different types of analyses that edgeR is capable of, including differential gene expression, differential exon usage, differential transcript expression, and time course analysis of RNA-seq data, as well as differential analysis of data from BS-seq, CRISPR-Cas9 and shRNA-seq genetic screens. The edgeR User’s Guide can be accessed by typing *edgeRUsersGuide()* at the R prompt. Questions related to edgeR analyses can be posted to the Bioconductor support site (https://support.bioconductor.org) and are generally answered promptly either by the authors or by other Bioconductor community members.

## Discussion

The edgeR package has been one of most widely used software tools for the statistical analysis of sequencing read counts over the past 15 years. The package develops and implements advanced statistical methods associated with generalized linear models, conditional likelihood and empirical Bayes for the analysis of count data. edgeR is used as an integrated analysis environment in its own right and is also used as an underlying engine by other packages that analyse specific technologies. At the time of writing (November 2024), 212 downstream Bioconductor packages depend on or suggest edgeR.

This article has summarized the design and capabilities of the edgeR package and has also described the history of the edgeR package over time. A detailed record of when each change was introduced is available from https://bioconductor.org/packages/release/bioc/news/edgeR/NEWS or by typing *news(package=“edgeR”)* at the R prompt. edgeR v4 introduces further new statistical ideas, improves computational efficiency and extends the range of applications of the package. Statistical innovations include modelling of fractional counts, a more refined modelling of the GLM deviances to achieve more accurate quasi-dispersion estimation in small count scenarios, and the idea of divided counts to extract out the overdispersion arising from transcript quantification. Implementation of low-level functions in C allows edgeR to handle larger datasets more efficiently. New data analyses include transcript-level differential expression, differential exon usage, differential transcript usage, differential methylation analysis, pseudo-bulk analysis of single-cell RNA-seq and hypothesis tests relative to a fold-change threshold. edgeR v4 also includes direct support for pathway analysis and gene set enrichment analysis.

## Data Availability

The edgeR package is freely available from https://-bioconductor.org/packages/edgeR. The source code can be browsed at https://code.bioconductor.org/-browse/edgeR or can be git cloned from /packages/edgeR. Additional documentation and datasets used in the edgeR User’s Guide are available from https://bioinf.wehi.edu.au/edgeR.

The example datasets shown in this article are publicly available as described in Materials and Methods. The TCGA data is available from https://tcga-data.nci.nih.gov. The RNA-seq data analysed by Chen *et al.* [10] and Fu *et al.* [35] is available as GEO series GSE60450. The RNA-seq data analysed by Fu *et al.* [36] is available as GEO series GSE118617.

## Acknowledgements

The authors thank all the colleagues who have been actively involved in the edgeR project over the years, including Mark Robinson, Davis McCarthy, Belinda Phipson, Matthew Ritchie, Zhiyin Dai, Oliver Voogd, Yifang Hu, Xiaobei Zhou, Jinming Cheng and Mengbo Li. The authors also appreciate all the suggestions and bug reports from the Bioconductor community as well as from edgeR users around the world.

